# Hypoxia couples growth and developmental timing by decoupling steroid synthesis and secretion

**DOI:** 10.64898/2026.01.12.699026

**Authors:** George P. Kapali, Alexander W. Shingleton

## Abstract

In almost all animals, low physiological levels of oxygen (hypoxia) reduce growth rate and adult body size. Despite the near ubiquity of this response, the systemic mechanisms that coordinate growth and development under hypoxia remain poorly understood. In *Drosophila*, hypoxia increases circulating levels of the steroid hormone ecdysone to inhibit insulin/IGF signaling and slow growth. At the same time, ecdysone biosynthesis is reduced to delay pupation and extend development. Traditionally, the secretion of lipid-soluble steroids is thought to be regulated at the level of biosynthesis. This therefore presents a paradox: how can a single environmental factor both increase ecdysone levels yet decrease ecdysone synthesis? Our data show that this paradox is resolved by the dual regulation of ecdysone at the levels of secretion and biosynthesis. We show that, in the short term, hypoxia increases basal levels of ecdysone via vesicle-mediated steroid hormone secretion, while in the long term, hypoxia decreases the expression of genes involved in ecdysone synthesis to delay the ecdysone peak that triggers pupation. We also present evidence that both ecdysone synthesis and secretion are regulated, in part, by NO-signaling. Collectively, our findings reveal that regulated steroid secretion as a critical and environmentally-responsive component of endocrine control, expanding our understanding of how animals integrate growth and developmental timing in response to acute and chronic environmental change.

**Author Summary:** Low oxygen levels (hypoxia) slow growth and development in almost all animals, but how changes in oxygen in the environment are sensed and turned into body-wide hormonal signals is still not well understood. Using *Drosophila*, we show that oxygen regulates the steroid hormone ecdysone to slow growth and development through two distinct mechanisms: fast hormone release and slower suppression of hormone synthesis. In the short-term, low oxygen triggers the active release of stored ecdysone from the gland that produces it, leading to a rise in circulating hormonal levels that slows body growth. In the longer-term, low oxygen reduces ecdysone production, delaying the hormone peak that initiates pupation and extending overall development. While it is widely accepted that fat-soluble steroids can pass freely, and passively, across membranes, these findings suggest that their active release allows a rapid response to environmental change. More broadly, our work suggests that environmental modulation of steroid secretion, alongside steroid synthesis, may represent a general strategy by which animals coordinate short- and long-term developmental responses to environmental stress.

## Introduction

Oxygen supports virtually all biological processes in most multicellular organisms; yet, its availability can vary widely across different developmental and ecological contexts. In almost all animals, low oxygen levels (hypoxia) during development reduce growth rate and final body size [1]. The evolutionary conservation of this response suggests that the mechanisms linking oxygen availability to growth are fundamental to animal development [2]. While we know much about the cellular response to hypoxia, which is mediated by the evolutionarily conserved hypoxia-inducible factor (HIF) signaling pathway [3], we know much less about the systemic mechanisms that coordinate hypoxic growth suppression across the whole organism. Identifying these mechanisms is essential for understanding how environmental oxygen shapes developmental trajectories and adult body size.

Morphologically and physiologically, the effects of developmental hypoxia closely mirror those of nutritional limitation. In *Drosophila melanogaster*, for example, hypoxia and dietary restriction both slow larval growth, extend development, and reduce adult body size [4]. Further, the effects of low oxygen and nutrition on body shape (i.e., relative trait size) are identical [5], which is not true for the impact of temperature on body size and shape [6]. Finally, adult body size is sensitive to both nutrition and oxygen, primarily during the third larval instar.

Collectively, these parallels suggest that oxygen and nutrient signals converge on shared endocrine pathways that regulate systemic growth, a hypothesis supported by several developmental studies[5,7–9] . In *Drosophila*, the principal mechanism regulating growth in response to nutrition is the insulin/insulin-like growth factor signaling (IIS) pathway [10]. The IIS pathway is highly evolutionarily conserved [11], and its activity is regulated in part through the nutrient-dependent release of insulin-like peptides (dILPs in *Drosophila*). Hypoxia reduces systemic IIS activity through at least two mechanisms. First, hypoxia acts via HIF signaling in the larval fat body to inhibit dILP release by the brain [7]. Second, hypoxia elevates basal levels of the steroid hormone ecdysone (**Figure 1a**), which suppresses IIS by inducing expression of Imp-L2, an insulin-binding protein that antagonizes insulin signaling [5].

**Figure 1.**
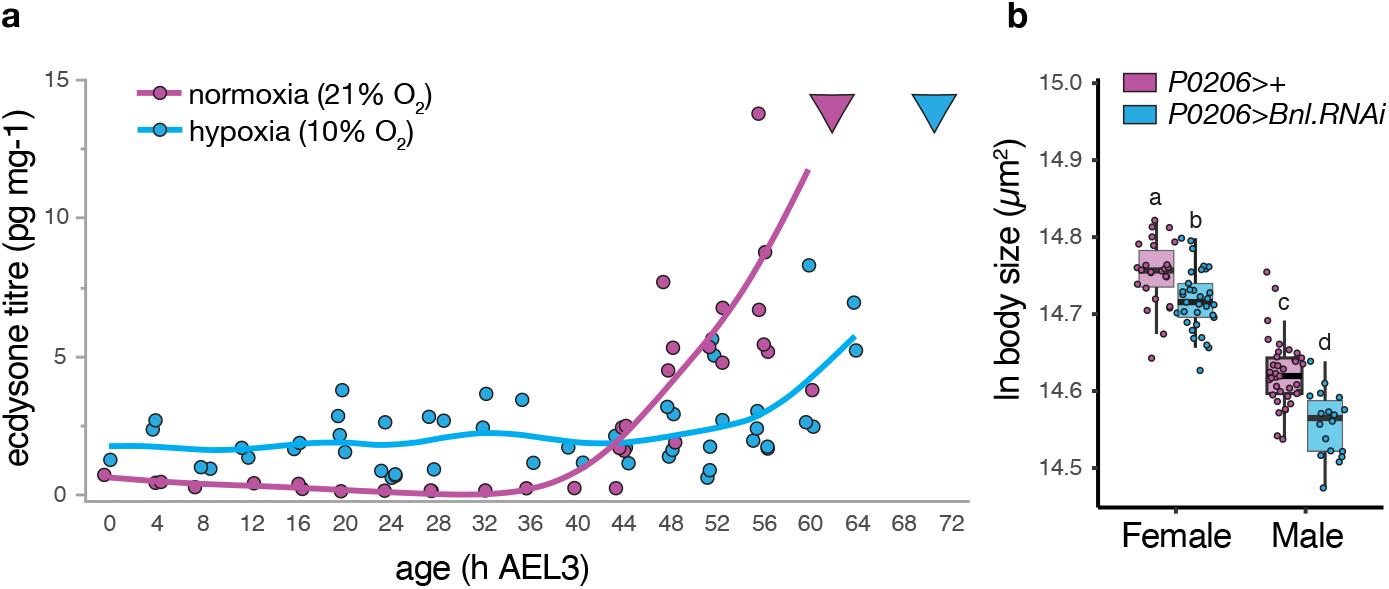
Hypoxia regulates ecdysone dynamics and growth. (A) Low oxygen (10%) increases basal levels of circulating ecdysone but delays the peak in ecdysone that initiates pupariation. Inverted triangles show mean age at pupariation. Lines are cubic splines. Data from [4]. (B) Disrupting tracheal development in the prothoracic gland (PG) through PG-specific knockdown of *branchless* reduces final body size (two-way ANOVA, F_genotype (1,111)_= 3.67, *P* < 0.0001 ). Error bars are 95% confidence intervals. Groups with different letters are significantly different at Šidák-corrected *P*<0.05.

While ecdysone is increasingly recognized as a regulator of growth rate in insects, it was first characterized as a regulator of developmental transitions, with pulses of ecdysone triggering larval-larval and larval-pupal molts [12]. In *Drosophila* and other holometabolous insects, the pulses of ecdysone that end larval growth and initiate pupation are associated with the attainment of a critical weight much earlier in the final instar [13]. In hypoxic *Drosophila* larvae, the critical weight is reduced such that metamorphosis is initiated in smaller larvae [14]. However, the period between attainment of critical weight and the ecdysone pulse that stops growth is extended [14] (**Figure 1a**). This delay is a consequence of reduced EGFR/ERK signaling in the prothoracic gland (PG: the primary site of ecdysone synthesis), which correspondingly reduces the expression of ecdysone synthesis genes (collectively called the Halloween genes) [8]. These findings, therefore, pose a paradox: if hypoxia suppresses ecdysone synthesis late in the third larval instar to delay pupariation, how does hypoxia simultaneously elevate basal ecdysone levels to reduce systemic growth early in the third larval instar?

In this study, we address this paradox by examining how low oxygen affects circulating ecdysone through its effects on the prothoracic gland. We confirm that the PG functions as an oxygen sensor and assess how hypoxia influences ecdysone synthesis, secretion, uptake, and degradation. Our findings show that hypoxia simultaneously slows growth and delays development by splitting ecdysone regulation into two distinct axes: secretion and synthesis. Canonically it is thought that steroid hormones diffuse freely across membranes [15–17]; however, increasing evidence from both insects and vertebrates demonstrates that steroid export and uptake are regulated, transporter-dependent processes [18–22] . We show that hypoxia enhances the secretion of pre-synthesized ecdysone early in the third larval instar to suppress growth, while later reducing ecdysone synthesis to delay the pulse that triggers metamorphosis. These results provide a new framework for understanding steroid hormone control, showing that their release, like their synthesis, can be actively modulated by environmental factors to shape developmental and physiological plasticity.

## Results

### The Prothoracic Gland is an Oxygen Sensor

Our previous study [5] demonstrated that ecdysone synthesis by the PG is necessary for growth suppression at 10% oxygen. An open question is whether the PG senses hypoxia directly or responds to signals from another oxygen-sensing tissue. Previous studies have shown that inhibiting tracheal development induces hypoxia in growing larvae and reduces adult body size. Tracheal development depends on the fibroblast growth factor (FGF)-like ligand Branchless. This extracellular signaling molecule is expressed by tissues in response to hypoxia gland (**Supplementary Figure S1a**), and acts as a guide to tracheal branching [23,24]. Knockdown of *Branchless*, whether ubiquitously [7] or in specific tissues [25], disrupts tracheal growth [26] and creates localized hypoxia [7]. To test whether limiting oxygen delivery to the prothoracic gland affects growth, we knocked down Branchless specifically in the PG using RNAi. We observed a significant reduction adult body size (**Figure 1b, Supplementary Figure S1b**). This phenotype is opposite to that expected if *Branchless* knockdown simply reduced PG growth, which is known to increase final body size [27] . Instead, these results suggest that impaired oxygen delivery to the PG is sufficient to suppress systemic growth, consistent with the PG directly sensing and responding to oxygen availability.

**Supplementary Figure S1.**
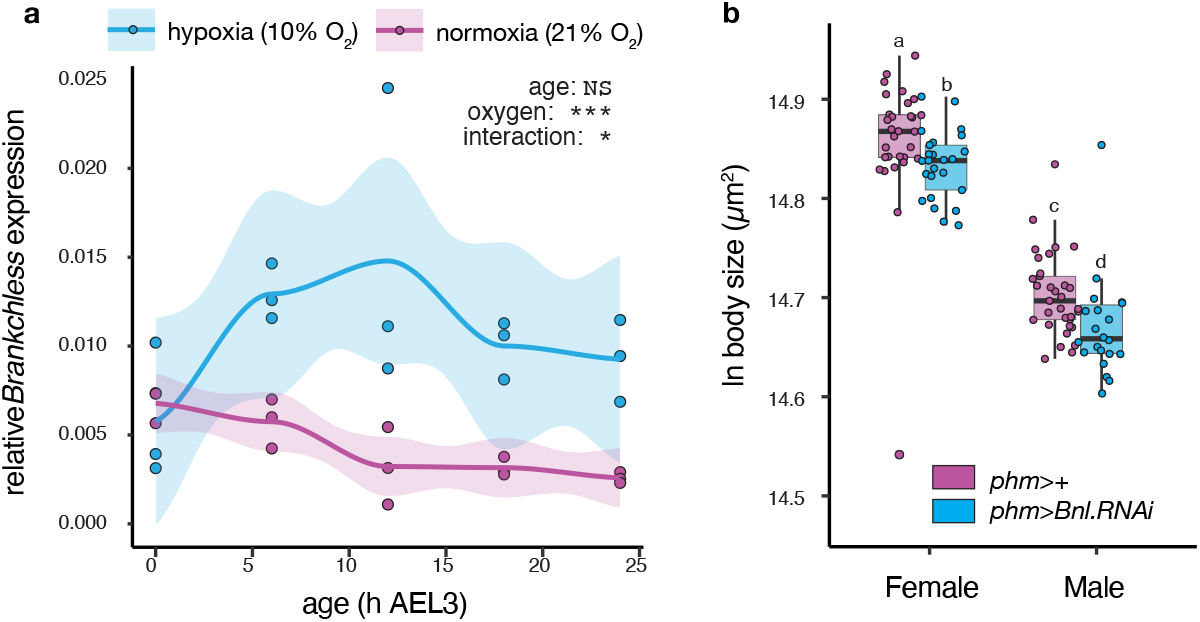
*Branchless* and hypoxia. (**a**) Expression of *Branchless* is up-regulated in response to hypoxia. Inset show result of two-way ANOVA: *expression* = *oxygen*\**age*. Line is cubic spline. (**b**) Disrupting tracheal development in the prothoracic gland through PG-specific knockdown of *Branchless* using the phm-GAL4 driver also reduces final body size (two-way ANOVA, F_genotype (1,107)_= 3.65, *P* < 0.0001 ). Error bars/confidence bands are 95% confidence intervals. Groups with different letters are significantly different at Šidák-corrected *P*<0.05.

We next tested whether hypoxia acts through HIF signaling in the prothoracic gland. When oxygen is plentiful, Hph (HIF prolyl hydroxylase) marks HIF-1α for degradation, there by inhibiting HIF-signaling [28] . If HIF-signaling in the PG reduces body size in hypoxia, then activating HIF-signaling through PG-specific knockdown of *Hph* in normoxia should have the same effect. Consistent with this prediction, PG-specific *Hph* knockdown reduced body size and delayed development, although in males only (**Supplementary Figure S2A & B**). However, PG-specific knockdown of *HIF-1α* produced similar reductions in body size and developmental delays, and did not rescue growth in hypoxia (**Supplementary Figure S2C**). Thus, neither activation nor inhibition of HIF signaling in the PG yields a consistent effect on growth or developmental timing. Previous studies have reported similarly mixed outcomes from manipulating HIF-signaling in the PG [9]. Together, these results indicate that the role of PG-specific HIF signaling in mediating the response to hypoxia is equivocal.

**Supplementary Figure S2.**
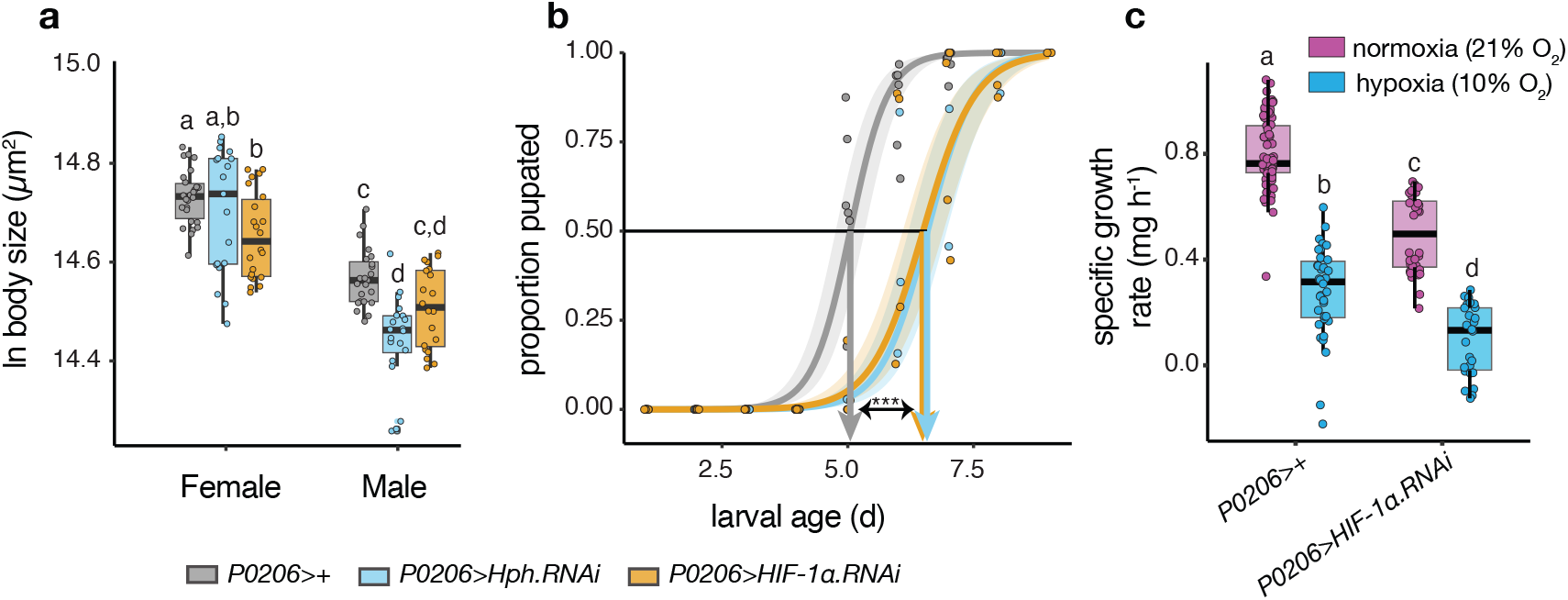
PG-specific HIF-signaling does not appear to regulate the response to hypoxia. (A) While activation of HIF-signaling in the PG (*P0206>Hph*.*RNAi*) decreases body size in normoxia, at least in males, so too does suppression of HIF-signaling *P0206>HIF-1α*.*RNAi*) (two-way ANOVA, *F*_*genotype*(1,124)_ =13.3142, *P*<0.0001 ) (B) Both activation and suppression of HIF-signaling in the PG delays development (GLMM, χ^2^ _genotype*age(2)_ = 23.284, *P*<0.0001). (**C**) Suppression of HIF-signaling through knockdown of *HIF-1α* does not rescue growth rate in hypoxia, but actually suppresses growth rate in both normoxia and hypoxia (two-way ANOVA, *F*_*genotype*\*oxygen(1,161)_ =5.9544, *P*=0.0158 ). Error bars/confidence bands are 95% confidence intervals. *** *P* < 0.001. Groups with different letters are significantly different at Šidák-corrected *P*<0.05.

### Hypoxia Does Not Elevate PTTH Signaling During Early Growth Suppression

The increase in basal levels of circulating ecdysone that suppress growth in hypoxia could be a response to an increase in activation of the PG by prothoracicotropic hormone (PTTH). This hormone is produced by the brain to stimulate PG growth and ecdysteroidgenesis [29] . and regulates the effect of environmental factors, such as crowding and nutrition, on body size and developmental timing [30] . To assay PTTH activity in response to hypoxia, we assayed the expression of *PTTH* and its receptor, *Torso*, in larvae that had been transferred to hypoxic conditions at ecdysis to the third instar. While hypoxia significantly elevated *PTTH* expression (**Figure 2a**), it did so only toward the end of the third larval instar (L3), after circulating ecdysone levels are elevated (**Figure 1a**). Hypoxia had no detectable effect on *Torso* expression (**Supplementary Figure S3a**).

**Figure 2.**
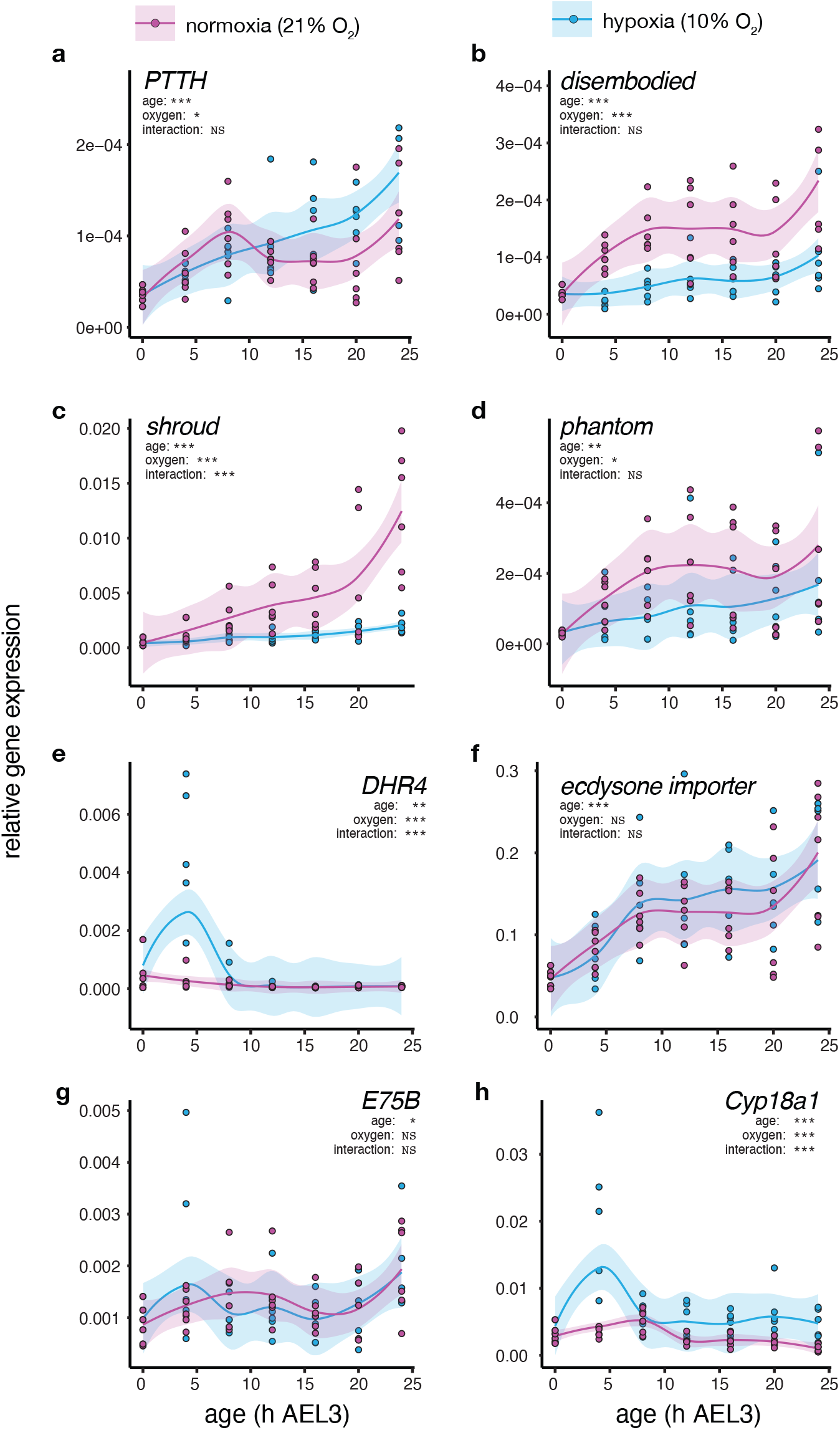
Effect of hypoxia on gene expression during the third larval instar. Larvae were transferred to hypoxia at ecdysis to L3. Charts are labeled by gene of interest. Inset show result of two-way ANOVA: *expression* = *oxygen*\**age*. Lines are cubic splines and confidence bands are 95% confidence intervals.

### Hypoxia Suppresses The Expression of Halloween Genes

The increase in basal levels of circulating ecdysone in hypoxia could be due to an increase in ecdysone synthesis. PTTH signaling directly targets the expression of ecdysone synthesis genes, collectively referred to as the Halloween genes, via the Ras-Raf-ERK pathway [31–35]. Genetic manipulation of Halloween gene expression in the PG disrupts ecdysone production, causing severe growth and timing defects that commonly prevent adult eclosion. When animals survive or when Halloween gene expression is altered indirectly via their upstream regulators, size at the end of larval development is strongly affected [33,36,37] . We therefore explored whether hypoxia increased the expression of these genes, focusing on the expression of *Neverland (Nvl), Shroud, Phantom, Disembodied*, and *Shadow* in early L3 reared under both hypoxia and normoxia. We found that hypoxia caused a general decrease in the expression of Halloween genes, delaying or eliminating the increase in Halloween gene expression that typically marks the end of larval development (**Figures 2b, c & d, Supplementary Figures S3b,c**). We also saw a discrete pulse of increased *DHR4* expression four hours after transfer to hypoxia at the L2/L3 molt (**Figure 2e**). DHR4 represses the transcription of multiple Halloween genes, including *Phantom* and *Disembodied*, and overexpressing *DHR4* suppresses the ecdysone pulses necessary for molting [32]. Collectively, these data are consistent with previous studies [8] demonstrating that hypoxia delays the ecdysone peak that drive pupariation by suppressing ecdysone biosynthesis.

**Supplementary Figure S3.**
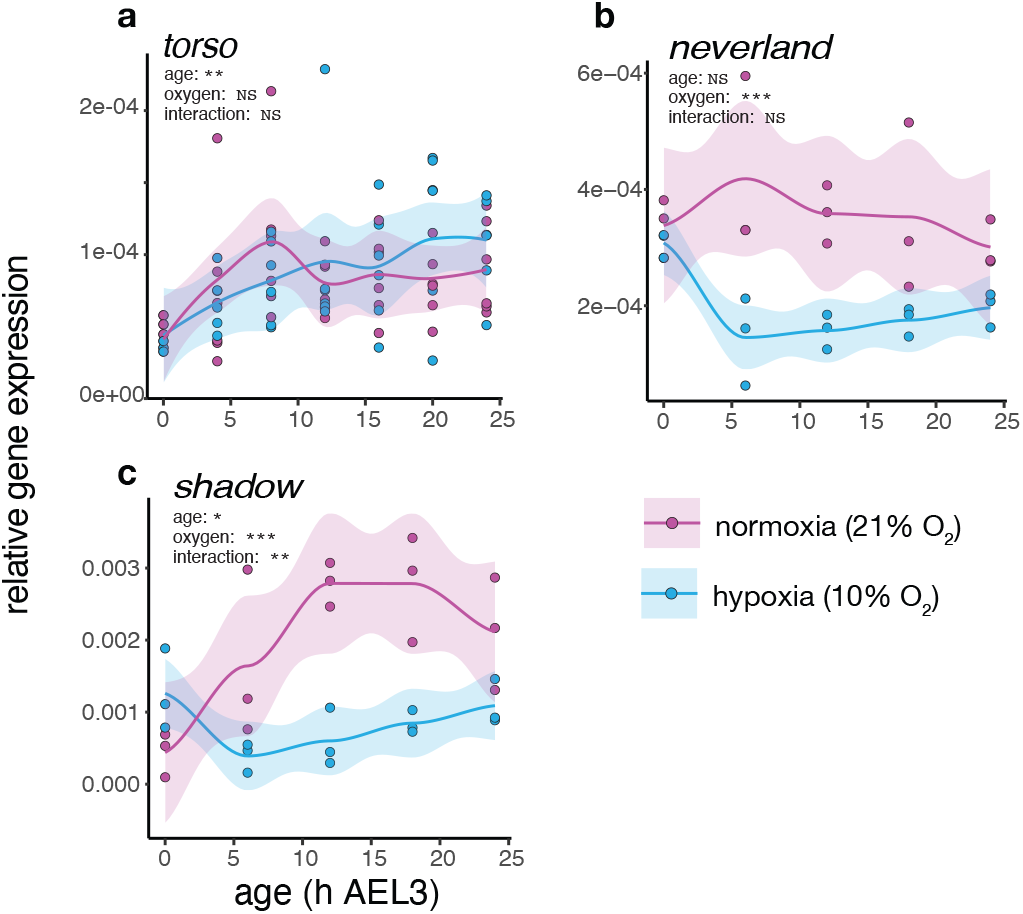
Effect of hypoxia on gene expression during the third larval instar. Larvae were transferred to hypoxia at ecdysis to L3. Charts are labeled by gene of interest. Inset show result of two-way ANOVA: *expression* = *oxygen*\**age*. Lines are cubic splines and confidence bands are 95% confidence intervals.

### Hypoxia Does Not Alter The Expression Of Ecdysone Importers Or Of Feedback-Regulator Genes

The increase in levels of circulating ecdysone in hypoxia could be a consequence of a decrease in the import of ecdysone into target tissues. We found, however, that hypoxia did not affect the expression of the ecdysone importer (*EcI*) (**Figure 2f**), a transmembrane protein that allows facilitate diffusion necessary for cellular uptake of ecdysone [22]. An increase in ecdysone could also reflect changes in the autoregulation of its own synthesis by the PG. This mechanism relies on signaling through the ecdysone signaling pathway, whereby ligand-bound ecdysone receptor Ecr and its co-receptor Usp activate DHR3 and E75. DHR3 then acts through FTZ-F1 to drive ecdysteroidgenesis, which is opposed by E75 (specifically the isoform E75B) [38,39]. Thus E75B serves as part of a negative feedback mechanism that regulates the timing of the ecdysone pulses at the end of larval development, but that may be altered in early L3 under hypoxia to elevate ecdysone synthesis. We found, however, no evidence that hypoxia affects the expression of *E75B* (**Figure 2g**).

### Hypoxia Does Not Decrease The Expression Of Ecdysone Degradation Genes

The elevated level of basal ecdysone observed in hypoxia may be due to a reduction in ecdysone degradation. We have previously shown that hypoxia does not down regulate the expression of *ecdysone oxidase* (*Eo*), which catalyzes the conversion of ecdysone to 3-dehydroecdysone (3DE), a crucial step in ecdysone inactivation [5]. We also measured the expression of *CYP18a1*, a key enzyme in ecdysteroid inactivation [40], and found that it is transiently elevated under hypoxic conditions (**Figure 2h**). Thus there is no evidence that the hypoxic-increase in basal ecdysone is a consequence of reduced ecdysone degradation.

### Hypoxia Transiently Increases The Expression Of The Ecdysone Transporter Atet

Collectively our data suggest that the increase in basal ecdysone levels that underlie hypoxic growth retardation are not due to changes in the synthesis, absorption or degradation of the steroid. We therefore tested whether hypoxia increased the transport of ecdysone out of the PG. Although ecdysone is a steroid and can, in principle, diffuse through cell membranes, recent evidence shows that its release from the PG involves vesicle exocytosis [21] . Ecdysone is loaded into vesicle by the ATP-binding cassette (ABC) transporter Atet (ABC transporter expressed in trachea), and exocytosis of the vesicles is stimulated by calcium-signaling sensed by Synaptotagmin1 (Syt1) in the vesicle membrane. To explore whether hypoxia induced the release of ecdysone from the PG, we assayed the expression of *Atet* in normoxic and hypoxic conditions. We found that *Atet* expression is transiently elevated four hours following transfer to hypoxia at ecdysis to L3 (**Figure 3A**). This increase in *Atet* expression coincides with the spike in *Imp-L2* expression that accounts for hypoxic reduction in growth rate [5]. To confirm this increase, we repeated our experiment and assayed *Atet* expression at ecdysis to L3 and four hours later, and found the same effect (**Figure 3B**). Thus, a compelling hypothesis is that hypoxia elevates basal levels of ecdysone by increasing its export from the PG.

**Figure 3.**
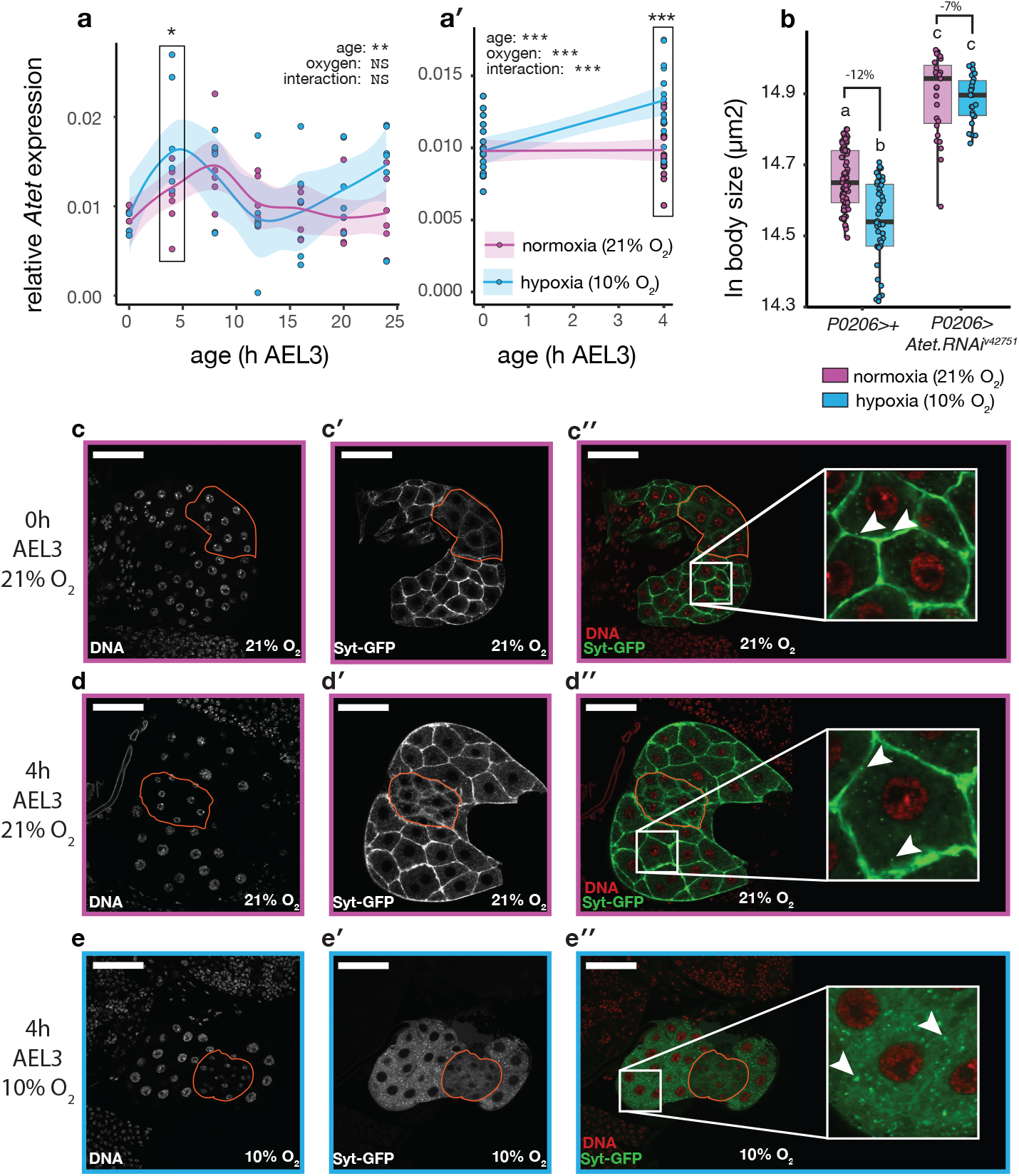
Hypoxia transiently increases the expression of *Atet* and induces exocytosis of Syt1 vesicles (**a, a′**) *Atet* expression for the first 24h (**a**) and 4h (**a′**) of L3. Insets show result of two-way ANOVA: expression = *oxygen*\**age*. Boxed points indicate hypoxic increase in *Atet* expression 4h AEL3 (t-test: *expression = oxygen*). Line is cubic spline and confidence bands are 95% confidence intervals. * *P*<0.05, *\*\*\* P* < 0.001. (**b**) PG-specific knockdown of *Atet* using RNAi eliminates the effects of hypoxia on body size (two-way ANOVA, *F*_*genotype*oxygen*(1,173)_ = 10.598, *P* = 0.0014). Groups with different letters are significantly different at Šidák-corrected *P*<0.05. (**c, d, e**) Representative image of *P0206>Syt-GFP* PG cells 0h AEL3 under normoxia (**c**), 4h AEL4 under normoxia (**d**) and 4h AEL4 under hypoxia (**e**). Under normoxia, Syt-GFP-tagged vesicles (arrowheads) aggregate at the cell membrane, while after 4 hours at hypoxia, this localization is lost, and the number of cytoplasmic vesicles increases. Orange line encloses the *corpora alatum*, which is not part of the PG. Scale bars are 10µm.

### Hypoxia Increases Vesicle-Mediated Secretion of Ecdysone from the PG

In their normal function, Syt1-positive secretory vesicles bind to the cell membrane to release their contents, before Syt1 is recycled back to form new vesicles [41] . Inhibiting calcium signaling in the PG, through knockdown of the *inositol1,4,5,-tris-phosphate receptor*(*IP3R*), results in accumulation of Syt1-positive secretory vesicles at the cell membrane [21] . If hypoxia induces the release of ecdysone through exocytosis, we should observe a decrease in Syt1-positive vesicles at the cell membrane as ecdysone is exocytosed into the hemolymph and Syt1 is recycled to form now secretory vesicles in the cytoplasm. To test this prediction we expressed eGFP-tagged Syt1 (*Syt-GFP*) in the PG to visualize the location ecdysone-containing vesicles in early L3 and their response to hypoxia. In normoxic larvae Syt1-positive vesicles accumulate at the membrane of PG cells both immediately after ecdysis to L3 and four hours later (**Figure 3 c & d**). In larvae maintained in hypoxia for four hours after ecdysis to L3 this accumulation is lost, and instead there is an increase in Syt1-positive vesicles in the cytoplasm (**Figure 3 e**). These observations are consistent with the hypothesis that hypoxia increases vesicle-mediated secretion of ecdysone from the PG.

### Atet Is Necessary for Growth Suppression Under Hypoxia

If hypoxia slows growth by increasing the Atet-mediated export of ecdysone from the PG, then blocking this export through knockdown of *Atet* should alleviate growth suppression. To test this we used RNAi to reduce the expression of *Atet* in the PG (*P0206*>*Atet*.*RNAi*^*50563*^) and found that it completely prevented growth suppression under hypoxia (**Figure 3 b**). This was accompanied by an overall increase in pupal size under both hypoxic and normoxic conditions, consistent with previous studies that show that inhibiting ecdysone release increases body size in *Drosophila* [27,29,42–44].

### NOS In The PG Is Necessary For Growth Suppression Under Hypoxia

Our results do not support the hypothesis that activation of HIF-signaling in the PG is responsible for growth suppression in response to hypoxia. Further, since the increase in basal ecdysone does not coincide with an increase in the expression of ecdysone biosynthetic genes, it seems unlikely that the response is due to signaling through either the insulin/IGF-signaling (IIS) or Ras/Raf/ERK-signaling pathway [33–35,45,46]. Several studies have linked the effects of hypoxia on physiology and growth to nitric oxide, a diffusible secondary messenger involved in various cellular functions [47–51]. Nitric oxide is generate by Nitric Oxide Synthase (NOS), which is known to disrupt the ecdysone’s negative feedback on its own synthesis, by preventing E75 from repressing DHR3 [38]. Furthermore, NO and calcium signaling are reciprocally linked: NOS activity is calcium-dependent [52,53], while NO feeds back to regulate intracellular calcium levels through several mechanisms [51,54]. A compelling hypothesis is that NOS is involved in the hypoxic suppression of growth by ecdysone.

To test this hypothesis, we first looked at the expression of *NOS* in normoxic and hypoxic larvae, and found that it was upregulated at low oxygen (**Figure 4a**). We next used RNAi to knock down *NOS* expression in the PG of normoxic and hypoxic larvae (*phm* > *NOS*.*RNAi*). We found that the loss of *NOS* partially, but significantly, alleviated the negative effect of hypoxia on body size (**Figure 4b**). These data confirm that *NOS* is necessary for complete growth suppression under hypoxia.

**Figure 4.**
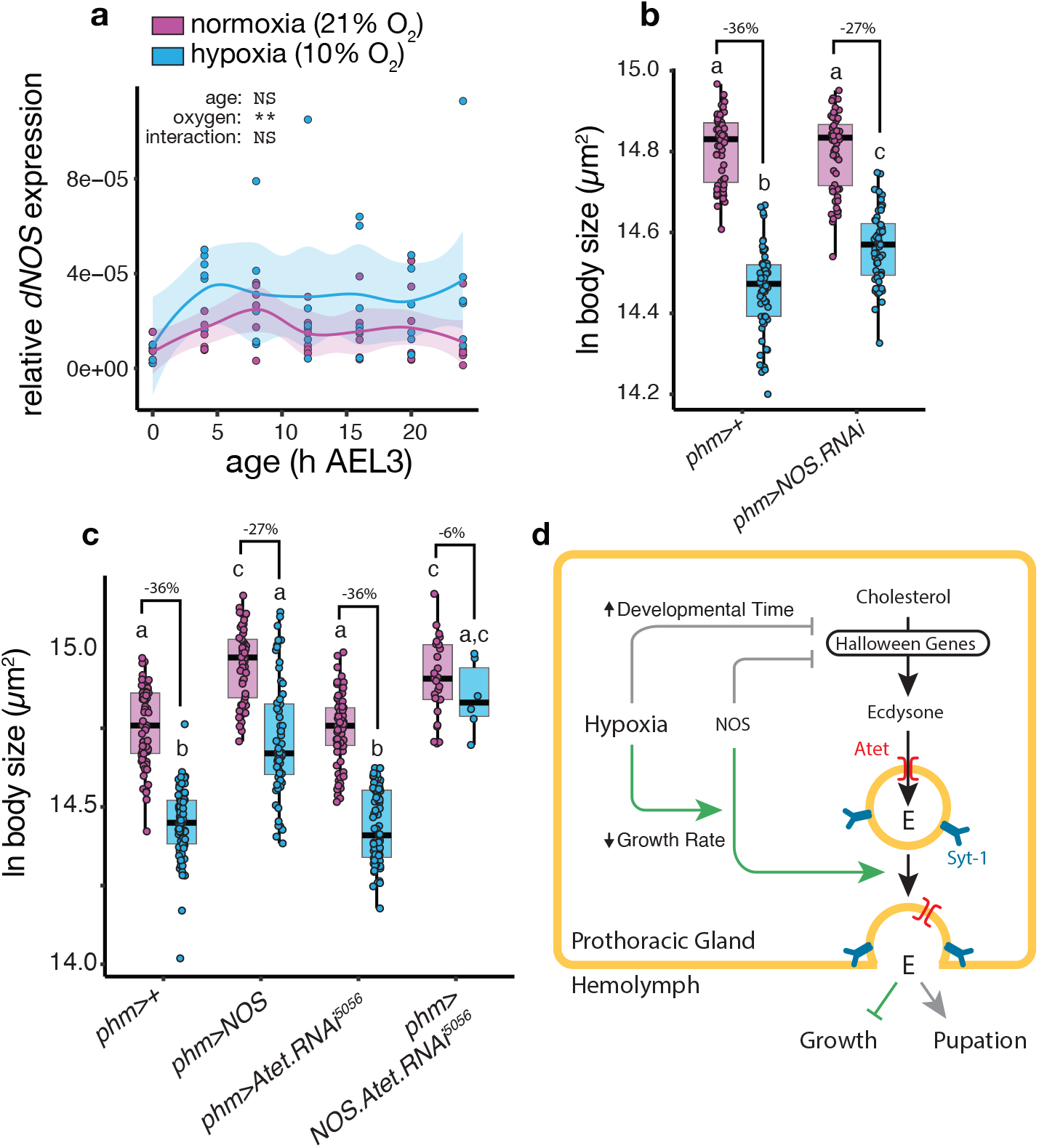
Hypoxia works through NOS to regulate body size. (**a**) Expression of *NOS* is upregulated in hypoxia. Inset shows result of two-way ANOVA: expression = *oxygen*\**age*. Lines are cubic splines and confidence bands are 95% confidence intervals. (**b**) PG-specific knockdown of *NOS* alleviates hypoxic size reduction (two-way ANOVA, *F*_*genotype*oxygen*(1,231)_ =17.0915, *P*<0.0001). (**c**) PG-specific overexpression of *NOS* increases body size and alleviates hypoxic size reduction (ANOVA, *F*_*genotype*oxygen*(1,270)_ =4.0554, *P*=0.04505). *NOS*-overexpression also sensitizes the PG to knockdown of *Atet* (two-way ANOVA, *F*_*genotype*oxygen*(1,189)_ =21.329, *P*<0.0001). Groups with different letters are significantly different at Šidák-corrected *P*<0.05. (**d**) A model of the effect of hypoxia on ecdysone synthesis and release, developmental time and growth rate.

If NOS is responsible for the increase in basal ecdysone levels that suppress growth in hypoxia, then upregulating its expression should genocopy the effects of low oxygen on growth and decrease body size in normoxia. Furthermore, if hypoxia acts through NOS signaling to increase the Atet-dependent export of ecdysone from the PG, the growth-inhibitory effects of increasing *NOS* expression in the PG should be mitigated by simultaneously knocking down *Atet* expression.

This, however, is not what we observed. Upregulating expression of *NOS* in the PG (*phm>NOS*) alone increased rather than decreased body size, again echoing the phenotypic effects of manipulations that inhibit ecdysone synthesis and release (**Figure 4c**). At the same time, the phenotypic plasticity of body size in response to oxygen was retained albeit slightly, but significantly, reduced (**Figure 4c**). We found that larvae where *NOS* and *Atet* expression were simultaneously increased and decreased, respectively (*phm< Atet. RNAi*^*v42751*^*;NOS*) died as second instars. We therefore used an ostensibly weaker *Atet* RNAi transgene (*Atet. RNAi*^*5056*^), which did not affect body size or oxic plasticity when expressed in the PG (**Figure 4c**). However, when co-expressed with NOS (*phm < Atet. RNAi*^*5056*^*;NOS*), this weaker knockdown completely prevented growth suppression under hypoxia while also increasing body size (**Figure 4bc**). Indeed, *phm>Atet. RNAi*^*5051*^*;NOS* larvae had the same body size and oxic plasticity as *phm*>*Atet. RNAi*^*v42751*^ larvae (**Figure 3b**). These data suggest that the co-expression of *NOS* renders the PG more sensitive to the knockdown of *Atet* in both normoxic and hypoxic conditions, placing *Atet* downstream of NOS in the regulation of growth and development by oxygen.

## Discussion

Hypoxia produces a puzzling combination of developmental effects in *Drosophila*. On the one hand, hypoxia reduces body size in part by elevating basal levels of circulating ecdysone, which suppresses insulin signaling via Imp-L2 and slows growth [4,5]. On the other hand hypoxia suppresses the expression of ecdysone biosynthetic genes, delaying the ecdysone peaks that trigger metamorphosis and thereby extending larval development [4,8]. Our findings resolve this paradox by demonstrating that ecdysone levels are regulated along two axes – biosynthesis and secretion – and suggests that environmental stress can decouple these regulatory processes. The increase in basal ecdysone results from enhanced Atet-dependent secretion of the hormone, rather than increased biosynthesis, supported by our direct observations of vesicles in the PG and by our finding that Atet is required for growth suppression under hypoxia (**Figure 3**). In contrast, the delay in the late ecdysone peaks reflects reduced ecdysteroidogenesis, supported by our observation that hypoxia lowers and delays the expression of the Halloween genes that encode ecdysteroidogenic enzymes (**Figure 2, Supplementary Figure S3**).

Classical models of steroid biology assume that steroids diffuse freely across membranes, such that circulating levels primarily reflect steroidogenic capacity [55–57]. Under this framework, environmental modulation of ecdysone is constrained by the rate at which biosynthetic gene expression and enzyme activity can change – a slow process, particularly under hypoxia, which suppresses transcription and metabolism. This is in contrast to peptide hormones, such as insulin, which can accumulate and be released rapidly via vesicle exocytosis [58]. The recent discovery of dedicated ecdysteroid exporters [21] and importers [22] suggests a similarly fast route for modulating ecdysone flux. Ecdysone-containing vesicles accumulate at the plasma membrane of prothoracic gland cells [21], and our data suggest that these vesicles rapidly exocytose their contents in response to hypyoxia. Starvation produces a similar dual effect in *Drosophila*, elevating basal levels of ecdysone to slow growth [59] while repressing biosynthetic gene expression [60], and may be regulated by the same mechanism. These patterns support a model (**Figure 4d**) in which environmental stresses differentially engage the secretory and biosynthetic arms of ecdysone regulation, allowing ecdysone to drive acute physiological responses while also reshaping longer-term developmental programs.

Our finding that restricting tracheal growth in the PG reduces body size suggests that the gland serves as an oxygen-responsive endocrine tissue. Nevertheless, the mechanism by which oxygen level is sensed by the PG to regulate the two arms of ecdysone synthesis and release remains unclear. The most obvious candidate for sensing hypoxia is the HIF-signaling pathway. Neither prior studies [9] nor our own data, however, provide clear evidence that HIF activity in the PG drives hypoxia-induced growth suppression. Instead, our data point to the involvement of NOS signaling, which is known to mediate the hypoxic response in mammals [51], and which has previously being implicated in the hypoxic response in insects [47,51].

Consistent with this, NOS appears to contribute to both axes of ecdysone regulation, albeit in different ways. Early in the third larval instar, NOS appears to be required, though not sufficient, for the oxygen-dependent release of ecdysone that elevates basal hormone levels and suppresses growth under hypoxia. Later in development, NOS becomes sufficient, though not required, to influence the timing of the ecdysone peaks that terminate growth and initiate metamorphosis, acting independently of oxygen. This context-dependence of NOS is mirrored in its role regulating the response to imaginal disc damage in *Drosophila*. Here a NOS-dependent mechanism in the PG represses ecdysone biosynthesis to slow and coordinate the growth of the undamaged discs, while a NOS-independent mechanism delays the PTTH-dependent peaks in ecdysone to extend developmental time and allow disc regeneration [61]. These observations may help explain the diverse, and sometimes conflicting, phenotypes reported after NOS manipulation in the PG [38,47,61]: NOS may act on both ecdysone synthesis and release, so that its effects depend on developmental stage, activation level, and stress context. Positioning NO/NOS within a two-axis regulatory model therefore helps clarify the divergent outcomes of NOS perturbation seen in *Drosophila* and other insects.

It is important to note that NOS signaling does not operate in isolation from other hypoxia-responsive pathways. In mammalian systems, NOS and HIF signaling are known to interact, with nitric oxide modulating HIF stability and activity, and HIF in turn influencing NO production[62]. Consistent with this, while HIF-signaling in the PG does not appear to drive hypoxia-induced growth suppression (**Supplementary Figure S2**), HIF is required for the regulation of developmental timing under hypoxia: loss of HIF activity has little effect on developmental timing in normoxia but substantially extends development in low oxygen [9]. In addition to local signaling within the PG, HIF-dependent signals from peripheral tissues, particularly the fat body, are also known to influence systemic growth regulation in hypoxia [63] and may act in parallel with, or upstream of, NOS-dependent mechanisms.

Together, these observations suggest that hypoxic regulation of growth and development emerges from the integration of NOS- and HIF-dependent signals across multiple tissues, rather than from a single, linear oxygen-sensing pathway.

In conclusion, our study provides a mechanism by which animals can rapidly adjust levels of circulating steroid in response to environmental stress, independent of steroid synthesis. Hypoxia induces the Atet-dependent release of ecdysone from the PG, accounting for the increase in basal levels of ecdysone that suppress body growth. Independently, hypoxia inhibits the expression of the Halloween genes, delaying the peaks in ecdysone that mark the end of larval development and retarding metamorphosis. In this way, ecdysone can regulate both the acute and chronic responses to environmental stress through the release and synthesis of ecdysone, respectively. Work in mammals has begun to elucidate the functions of steroid exporters, including ABC transporters [64–66], and importers such as megalin and OATPs [18,67], yet how these systems shape the dynamics of steroid availability is still unclear [68]. Our results suggest that, in insects, transporter activity provides a fast regulatory route for modifying circulating ecdysone levels, revealing a layer of control that parallels the vesicular regulation of peptide hormones.

## Materials and Methods

### Drosophila Stocks and Maintenance

We used the following fly strains: the isogenic RNAi control (VDRC 60,000), *UAS Atet RNAi*^*v42751*^ (VDRC 42751), *phm-GAL4* (a gift from Michael O’Connor), *P0206-Gal4* (a gift from Christen Mirth), *UAS-Syt-GFP* (BDSC 6925), *UAS-Atet RNAi*^*50563*^ (BDSC 50563), and *UAS-Bnl RNAi* (BDSC 34572). All constructs were backcrossed into VDRC 60,000, for five generations prior to analysis. Flies were reared at low density (50-100 larvae per vial), 25°C, 21% O_2_ in constant light on a sucrose-yeast diet containing 13 g of carrageenan, 100 g of yeast extract, and 50 g of sucrose in 1000 ml of water. To prevent bacterial and fungal growth, we added 3% Nipagin and 0.3% (v/v) propionic acid to the cooled mixtures.

### Body Size Measurement

All larvae were reared under standard conditions (25°C, 21% O_2_) and staged into 2h cohorts at ecdysis to the third instar before being maintained at 25°C, 21% O_2_ (normoxic flies) or moved to 25°C, 10% O_2_ (hypoxic flies) for the remainder of their development. We used pupal case size as a measure of body size [69] . We collected digital images of the pupal cases and measured the area of the pupal case when viewed from the dorsal aspect using ImageJ. Measurements were log transformed prior to analysis.

### Growth Rate Measurements

*P0206>+* and *P0206>HIF-1α*.*RNAi* larvae were reared under standard conditions (25°C, 21% O_2_) and massed at ecdysis to L3 (72h after oviposition) and 24h later (96h after oviposition. Specific growth rate (mg d^-1^) was calculated as 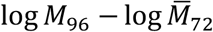, where *M*_*x*_ is mass at age *x*.

### Quantitative PCR (qPCR)

VDRC *60,000* larvae were reared under standard conditions (25°C, 21% O_2_) and staged into 2h cohorts at ecdysis to the third instar before being maintained at 25°C, 21% O_2_ (normoxic flies) or moved to 25°C, 10% O_2_ (hypoxic flies). Larvae were subsequently collected every four or six hours for 24 hours and preserved in groups of ten in RNAlater (Thermo Fischer Scientific). We extracted the RNA using Trizol (Invitrogen), treated it with DNase I (Thermo Fischer Scientific), quantified it with a NanoDrop One (Thermo Fischer Scientific) and reverse transcribed it with a High-Capacity cDNA Reverse Transcrip9on Kit (Applied Biosystems). Quantitative RT–PCR was conducted using PowerUp SYBR Green Master Mix (Applied Biosystems) and measured on a QuantStudio 3 Real-Time PCR system (Applied Biosystems). We calculated mRNA abundance for three biological replicates of ten larvae, using a standard curve and normalizing against the expression of ribosomal protein 49 (*RP49*), as described previously [5]. Standard curves were generated using seven serial dilutions of total RNA extracted from the isogenic control line (60,000): 5x 1st instar larvae, 5x 2nd instar larvae, 3rd instar larvae (male), 5x pupae (male), 5x adult flies (male). Primer sequences used in the study are listed in **Supplementary Table S1**.

### Syt-GFP-Positive Vesicle Localization

*P0206< Syt-GFP* larvae were reared as described above, then dissected either at ecdysis to the third larval instar or four hours later following maintenance under normoxia (21% O2) or hypoxia (10% O2). We separated the anterior third of the larval body and inverted the head in ice-cold PBS before fixing in 4% PFA for 30 minutes. We then dissected the fixed brain-ring gland complex from the head, removing the esophagus and gut, which the ring-gland encircles. The brain-ring gland complex was then transferred to 0.3% PBT and washed for 1 hour, before being mounted in Vectashield Antifade Mounting Media with DAPI. The PGs and surround tissue were then imaged under a confocal microscope.

### Statistical Analyses

All statistical analyses were conducted in *R* [70]. All the data and the R-scripts used to analyze them are provided on Dryad (DOI:10.5061/dryad.w9ghx3g4f). The effects of genotype on the response of body size and specific growth rate to hypoxia were tested using two-way ANOVAs (*body size/specific growth rate = genotype*oxygen*), with *post hoc* Šidák-corrected comparisons. The effect of genotype on developmental time was tested using a binomial GLMM (*pupation* = *age*genotype* + *vial*), where *vial* was treated as a random factor, with *post hoc* Tukey-corrected pairwise comparisons between genotypes. The effect of oxygen level on gene expression was tested using two-way ANOVAs (*expression = age*oxygen*), where *age* is a categorical variable, with *post hoc* Šidák-corrected comparisons within specific ages.

## Acknowledgments

This work was supported by grants from the National Science Foundation (US): IOS-1952385 and IOS-1406547, to AWS.

---

**Supplementary Table S1:**
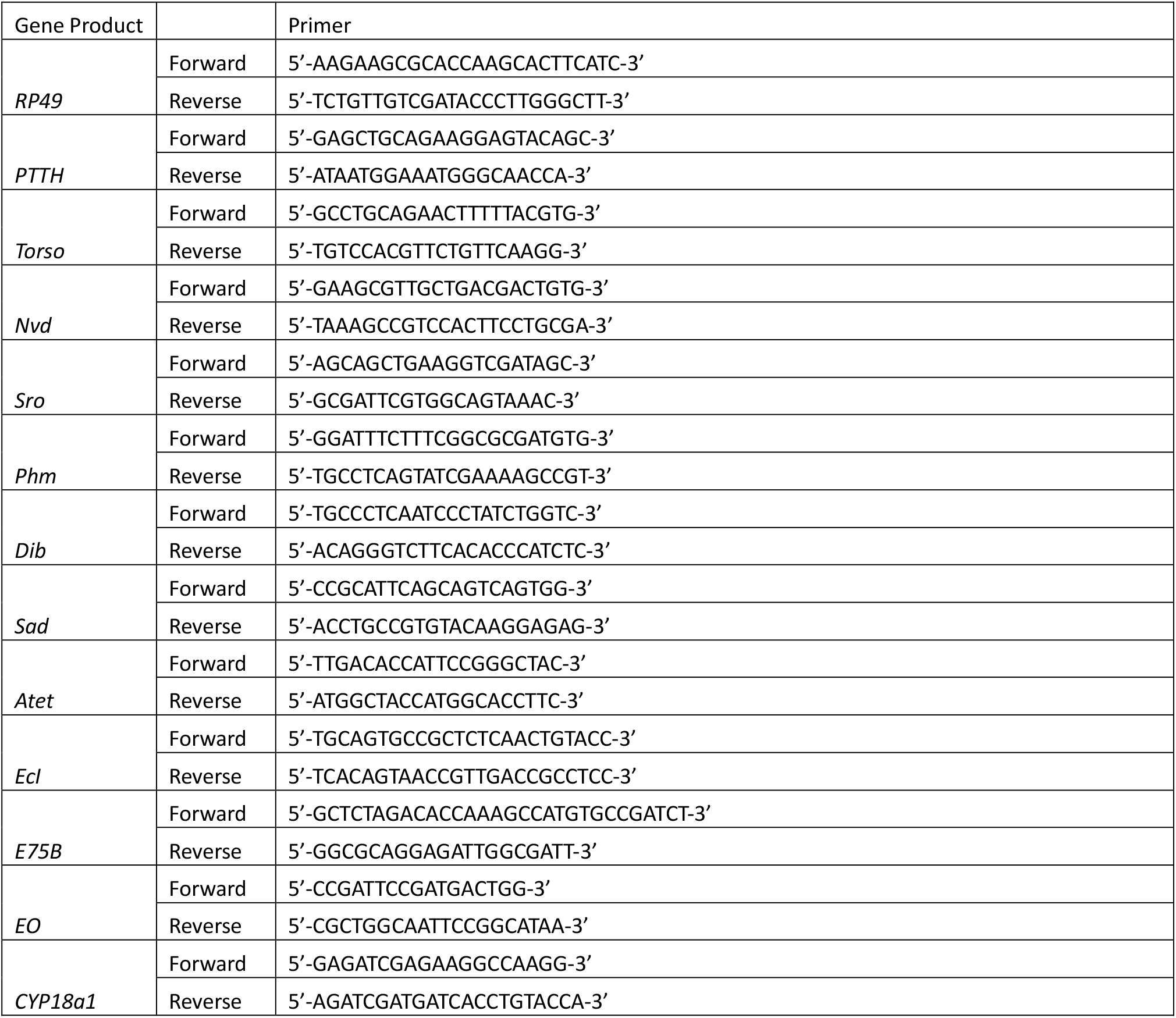
Oligonucleotide sequences used for qPCR.

## References

1. Harrison JF, Shingleton AW, Callier V. Stunted by Developing in Hypoxia: Linking Comparative and Model Organism Studies. Physiol Biochem Zool. 2015;88: 455 470. doi:10.1086/682216

2. Hampton-Smith RJ, Peet DJ. From Polyps to People. Ann N York Acad Sci. 2009;1177: 19–29. doi:10.1111/j.1749-6632.2009.05035.x

3. Loenarz C, Coleman ML, Boleininger A, Schierwater B, Holland PWH, Ratcliffe PJ, et al. The hypoxia-inducible transcription factor pathway regulates oxygen sensing in the simplest animal, Trichoplax adhaerens. EMBO Rep. 2011;12: 63–70. doi:10.1038/embor.2010.170

4. Callier V, Shingleton AW, Brent CS, Ghosh SM, Kim J, Harrison JF. The role of reduced oxygen in the developmental physiology of growth and metamorphosis initiation in Drosophila melanogaster. J Exp Biol. 2013;216: 4334 4340. doi:10.1242/jeb.093120

5. Kapali GP, Callier V, Gascoigne SJL, Harrison JF, Shingleton AW. The steroid hormone ecdysone regulates growth rate in response to oxygen availability. Sci Rep-uk. 2022;12: 4730. doi:10.1038/s41598-022-08563-9

6. Shingleton AW, Estep CM, Driscoll MV, Dworkin I. Many ways to be small: different environmental regulators of size generate distinct scaling relationships in Drosophila melanogaster. Proc Royal Soc B Biological Sci. 2009;276: 2625–2633. doi:10.1098/rspb.2008.1796

7. Texada MJ, Jørgensen AF, Christensen CF, Koyama T, Malita A, Smith DK, et al. A fat-tissue sensor couples growth to oxygen availability by remotely controlling insulin secretion. Nat Commun. 2019;10: 1955. doi:10.1038/s41467-019-09943-y

8. Turingan MJ, Li T, Wright J, Sharma A, Ding K, Khan S, et al. Hypoxia delays steroid-induced developmental maturation in Drosophila by suppressing EGF signaling. PLOS Genet. 2024;20: e1011232. doi:10.1371/journal.pgen.1011232

9. Campbell JB, Shingleton AW, Greenlee KJ, Gray AE, Smith HC, Callier V, et al. HIF signaling in the prothoracic gland regulates growth and development in hypoxia but not normoxia in Drosophila. J Exp Biol. 2024;227. doi:10.1242/jeb.247697

10. Brogiolo W, Stocker H, Ikeya T, Rintelen F, Fernandez R, Hafen E. An evolutionarily conserved function of the Drosophila insulin receptor and insulin-like peptides in growth control. Curr Biol. 2001;11: 213 221. doi:10.1016/s0960-9822(01)00068-9

11. Chan SJ, Steiner DF. Insulin Through the Ages: Phylogeny of a Growth Promoting and Metabolic Regulatory Hormone1. Am Zoöl. 2000;40: 213–222. doi:10.1093/icb/40.2.213

12. Riddiford LM, Cherbas P, Truman JW. Ecdysone receptors and their biological actions. Vitamins Hormones. 2000;60: 1 73. doi:10.1016/s0083-6729(00)60016-x

13. Nijhout HF, Riddiford LM, Mirth C, Shingleton AW, Suzuki Y, Callier V. The developmental control of size in insects. Wiley Interdiscip Rev Dev Biology. 2014;3: 113 134. doi:10.1002/wdev.124

14. Callier V, Hand SC, Campbell JB, Biddulph T, Harrison JF. Developmental changes in hypoxic exposure and responses to anoxia in Drosophila melanogaster. J Exp Biol. 2015;218: 2927–2934. doi:10.1242/jeb.125849

15. Alberts B, Johnson A, Lewis J, Morgan D, M. Raff, K. Roberts, et al. Molecular Biology of the Cell. Sixth Edition. Garland Publishing Science; 2015.

16. Evans RM. The steroid and thyroid hormone receptor superfamily. Science. 1988;240: 889–95. doi:10.1126/science.3283939

17. Urry LA, Cain ML, Wasserman SA, Minorsky PV, Reece JB. Campbell Biology. 11th ed. New York: Pearson; 2017.

18. Hammes A, Andreassen TK, Spoelgen R, Raila J, Hubner N, Schulz H, et al. Role of Endocytosis in Cellular Uptake of Sex Steroids. Cell. 2005;122: 751–762. doi:10.1016/j.cell.2005.06.032

19. Willnow TE, Nykjaer A. Cellular uptake of steroid carrier proteins—Mechanisms and implications. Mol Cell Endocrinol. 2010;316: 93–102. doi:10.1016/j.mce.2009.07.021

20. Simon E, Bonche R, Maarouf Y, Nawrot-Esposito M-P, Romero NM. Filopodia are essential for steroid release. Nat Commun. 2025;16: 5501. doi:10.1038/s41467-025-60579-7

21. Yamanaka N, Marqués G, O’Connor MB. Vesicle-Mediated Steroid Hormone Secretion in Drosophila melanogaster. Cell. 2015;163: 907 919. doi:10.1016/j.cell.2015.10.022

22. Okamoto N, Viswanatha R, Bittar R, Li Z, Haga-Yamanaka S, Perrimon N, et al. A Membrane Transporter Is Required for Steroid Hormone Uptake in Drosophila. Dev Cell. 2018;47: 294-305.e7. doi:10.1016/j.devcel.2018.09.012

23. Sutherland D, Samakovlis C, Krasnow MA. branchless Encodes a Drosophila FGF Homolog That Controls Tracheal Cell Migration and the Pattern of Branching. Cell. 1996;87: 1091–1101. doi:10.1016/s0092-8674(00)81803-6

24. Centanin L, Dekanty A, Romero N, Irisarri M, Gorr TA, Wappner P. Cell Autonomy of HIF Effects in Drosophila: Tracheal Cells Sense Hypoxia and Induce Terminal Branch Sprouting. Dev Cell. 2008;14: 547–558. doi:10.1016/j.devcel.2008.01.020

25. Sauerwald J, Backer W, Matzat T, Schnorrer F, Luschnig S. Matrix metalloproteinase 1 modulates invasive behavior of tracheal branches during entry into Drosophila flight muscles. eLife. 2019;8: e48857. doi:10.7554/elife.48857

26. Tamamouna V, Rahman MM, Petersson M, Charalambous I, Kux K, Mainor H, et al. Remodelling of oxygen-transporting tracheoles drives intestinal regeneration and tumorigenesis in Drosophila. Nat Cell Biol. 2021;23: 497–510. doi:10.1038/s41556-021-00674-1

27. Mirth C, Truman J, Riddiford L. The Role of the Prothoracic Gland in Determining Critical Weight for Metamorphosis in. Curr Biol. 2005;15: 1796 1807. doi:10.1016/j.cub.2005.09.017

28. Lavista-Llanos S, Centanin L, Irisarri M, Russo DM, Gleadle JM, Bocca SN, et al. Control of the hypoxic response in Drosophila melanogaster by the basic helix-loop-helix PAS protein similar. Molecular and Cellular Biology. 2002;22: 6842 6853.

29. Mcbrayer Z, Ono H, Shimell M, Parvy J-P, Beckstead RB, Warren JT, et al. Prothoracicotropic Hormone Regulates Developmental Timing and Body Size in Drosophila. Dev Cell. 2007;13: 857 871. doi:10.1016/j.devcel.2007.11.003

30. Shimell M, Pan X, Martin FA, Ghosh AC, Leopold P, O’Connor MB, et al. Prothoracicotropic hormone modulates environmental adaptive plasticity through the control of developmental timing. Development. 2018;145: dev159699. doi:10.1242/dev.159699

31. Rewitz KF, Yamanaka N, Gilbert LI, O’connor MB. The Insect Neuropeptide PTTH Activates Receptor Tyrosine Kinase Torso to Initiate Metamorphosis. Science. 2009;326: 1403 1405. doi:10.1126/science.1176450

32. Ou Q, Magico A, King-Jones K. Nuclear receptor DHR4 controls the timing of steroid hormone pulses during Drosophila development. Plos Biol. 2011;9: e1001160. doi:10.1371/journal.pbio.1001160

33. Danielsen ET, Moeller ME, Dorry E, Komura-Kawa T, Fujimoto Y, Troelsen JT, et al. Transcriptional Control of Steroid Biosynthesis Genes in the Drosophila Prothoracic Gland by Ventral Veins Lacking and Knirps. PLoS Genet. 2014;10: e1004343. doi:10.1371/journal.pgen.1004343

34. Cruz J, Martín D, Franch-Marro X. Egfr Signaling Is a Major Regulator of Ecdysone Biosynthesis in the Drosophila Prothoracic Gland. Curr Biol. 2020;30: 1547-1554.e4. doi:10.1016/j.cub.2020.01.092

35. Kannangara JR, Mirth CK, Warr CG. Regulation of ecdysone production in Drosophila by neuropeptides and peptide hormones. Open Biol. 2021;11: 200373. doi:10.1098/rsob.200373

36. Enya S, Ameku T, Igarashi F, Iga M, Kataoka H, Shinoda T, et al. A Halloween gene noppera-bo encodes a glutathione S-transferase essential for ecdysteroid biosynthesis via regulating the behaviour of cholesterol in Drosophila. Sci Rep. 2014;4: 6586. doi:10.1038/srep06586

37. Ou Q, Zeng J, Yamanaka N, Brakken-Thal C, O’connor MB, King-Jones K. The Insect Prothoracic Gland as a Model for Steroid Hormone Biosynthesis and Regulation. Cell Reports. 2016;16: 247 262. doi:10.1016/j.celrep.2016.05.053

38. Cáceres L, Necakov AS, Schwartz C, Kimber S, Roberts IJH, Krause HM. Nitric oxide coordinates metabolism, growth, and development via the nuclear receptor E75. Gene Dev. 2011;25: 1476–1485. doi:10.1101/gad.2064111

39. White KP, Hurban P, Watanabe T, Hogness DS. Coordination of Drosophila Metamorphosis by Two Ecdysone-Induced Nuclear Receptors. Science. 1997;276: 114–117. doi:10.1126/science.276.5309.114

40. Guittard E, Blais C, Maria A, Parvy J-P, Pasricha S, Lumb C, et al. CYP18A1, a key enzyme of Drosophila steroid hormone inactivation, is essential for metamorphosis. Dev Biol. 2011;349: 35–45. doi:10.1016/j.ydbio.2010.09.023

41. Small C, Harper C, Jiang A, Kontaxi C, Pronot M, Yak N, et al. SV2A controls the surface nanoclustering and endocytic recruitment of Syt1 during synaptic vesicle recycling. J Neurochem. 2024;168: 3188–3208. doi:10.1111/jnc.16186

42. Caldwell PE, Walkiewicz M, Stern M. Ras activity in the Drosophila prothoracic gland regulates body size and developmental rate via ecdysone release. Curr Biol. 2005;15: 1785 1795. doi:10.1016/j.cub.2005.09.011

43. Pan X, O’Connor MB. Coordination among multiple receptor tyrosine kinase signals controls Drosophila developmental timing and body size. Cell Reports. 2021;36: 109644–109644. doi:10.1016/j.celrep.2021.109644

44. Ghosh A, Mcbrayer Z, O’connor MB. The Drosophila gap gene giant regulates ecdysone production through specification of the PTTH-producing neurons. Dev Biol. 2010;347: 271 278. doi:10.1016/j.ydbio.2010.08.011

45. Boulan L, Milán M, Léopold P. The Systemic Control of Growth. Csh Perspect Biol. 2015;7: a019117. doi:10.1101/cshperspect.a019117

46. Layalle S, Arquier N, Léopold P. The TOR Pathway Couples Nutrition and Developmental Timing in Drosophila. Dev Cell. 2008;15: 568 577. doi:10.1016/j.devcel.2008.08.003

47. DeLalio LJ, Dion SM, Bootes AM, Smith WA. Direct effects of hypoxia and nitric oxide on ecdysone secretion by insect prothoracic glands. J Insect Physiol. 2015;76: 56 66. doi:10.1016/j.jinsphys.2015.02.009

48. Jeong S. Function and regulation of nitric oxide signaling in Drosophila. Mol Cells. 2024;47: 100006. doi:10.1016/j.mocell.2023.12.004

49. DiGregorio PJ, Ubersax JA, O’Farrell PH. Hypoxia and Nitric Oxide Induce a Rapid, Reversible Cell Cycle Arrest of the Drosophila Syncytial Divisions*. J Biol Chem. 2001;276: 1930–1937. doi:10.1074/jbc.m003911200

50. Wingrove J, Ofarrell P. Nitric Oxide Contributes to Behavioral, Cellular, and Developmental Responses to Low Oxygen in. Cell. 1999;98: 105 114. doi:10.1016/s0092-8674(00)80610-8

51. Dijkers PF, O’Farrell PH. Dissection of a Hypoxia-induced, Nitric Oxide–mediated Signaling Cascade. Mol Biol Cell. 2009;20: 4083–4090. doi:10.1091/mbc.e09-05-0362

52. Esplugues JV. NO as a signalling molecule in the nervous system. Br J Pharmacol. 2002;135: 1079–1095. doi:10.1038/sj.bjp.0704569

53. Regulski M, Tully T. Molecular and biochemical characterization of dNOS: a Drosophila Ca2+/calmodulin-dependent nitric oxide synthase. Proc Natl Acad Sci. 1995;92: 9072–9076. doi:10.1073/pnas.92.20.9072

54. Clementi E. Role of Nitric Oxide and Its Intracellular Signalling Pathways in the Control of Ca2 Homeostasis. Biochem Pharmacol. 1998;55: 713–718. doi:10.1016/s0006-2952(97)00375-4

55. Oren I, Fleishman SJ, Kessel A, Ben-Tal N. Free Diffusion of Steroid Hormones Across Biomembranes: A Simplex Search with Implicit Solvent Model Calculations. Biophys J. 2004;87: 768–779. doi:10.1529/biophysj.103.035527

56. Rao GS. Mode of entry of steroid and thyroid hormones into cells. Mol Cell Endocrinol. 1981;21: 97–108. doi:10.1016/0303-7207(81)90047-2

57. Mendel CM. The Free Hormone Hypothesis: A Physiologically Based Mathematical Model*. Endocr Rev. 1989;10: 232–274. doi:10.1210/edrv-10-3-232

58. Ma L, Bindokas VP, Kuznetsov A, Rhodes C, Hays L, Edwardson JM, et al. Direct imaging shows that insulin granule exocytosis occurs by complete vesicle fusion. Proc Natl Acad Sci. 2004;101: 9266–9271. doi:10.1073/pnas.0403201101

59. Lee GJ, Han G, Yun HM, Lim JJ, Noh S, Lee J, et al. Steroid signaling mediates nutritional regulation of juvenile body growth via IGF-binding protein in Drosophila. Proc National Acad Sci. 2018;115: 201718834. doi:10.1073/pnas.1718834115

60. Koyama T, Rodrigues MA, Athanasiadis A, Shingleton AW, Mirth CK. Nutritional control of body size through FoxO-Ultraspiracle mediated ecdysone biosynthesis. Elife. 2014;3. doi:10.7554/elife.03091

61. Jaszczak JS, Wolpe JB, Dao AQ, Halme A. Nitric Oxide Synthase Regulates Growth Coordination During Drosophila melanogaster Imaginal Disc Regeneration. Genetics. 2015;200: 1219 1228. doi:10.1534/genetics.115.178053

62. Hendrickson MD, Poyton RO. Crosstalk between nitric oxide and hypoxia-inducible factor signaling pathways: an update. Res Rep Biochem. 2015;5: 147–161. doi:10.2147/rrbc.s58280

63. Texada MJ, Joergensen AF, Smith DK, Marple DFM, Danielsen ET, Petersen SK, et al. A fat-tissue sensor couples growth to oxygen availability by remotely controlling insulin secretion. Biorxiv. 2018; 348334. doi:10.1101/348334

64. Kerr ID, Haider AJ, Gelissen IC. The ABCG family of membrane-associated transporters: you don’t have to be big to be mighty. Br J Pharmacol. 2011;164: 1767–1779. doi:10.1111/j.1476-5381.2010.01177.x

65. Grube M, Hagen P, Jedlitschky G. Neurosteroid Transport in the Brain: Role of ABC and SLC Transporters. Front Pharmacol. 2018;9: 354. doi:10.3389/fphar.2018.00354

66. Devine K, Villalobos E, Kyle CJ, Andrew R, Reynolds RM, Stimson RH, et al. The ATP-binding cassette proteins ABCB1 and ABCC1 as modulators of glucocorticoid action. Nat Rev Endocrinol. 2023;19: 112–124. doi:10.1038/s41574-022-00745-9

67. Birn H, Zhai X, Holm J, Hansen SI, Jacobsen C, Christensen EI, et al. Megalin binds and mediates cellular internalization of folate binding protein. FEBS J. 2005;272: 4423–4430. doi:10.1111/j.1742-4658.2005.04857.x

68. Schweizer U, Braun D, Forrest D. The Ins and Outs of Steroid Hormone Transport Across the Plasma Membrane: Insight From an Insect. Endocrinology. 2018;160: 339–340. doi:10.1210/en.2018-01034

69. Stillwell RC, Dworkin I, Shingleton AW, Frankino WA. Experimental manipulation of body size to estimate morphological scaling relationships in Drosophila. J Vis Exp Jove. 2011; e3162.e3162. doi:10.3791/3162

70. R_Core_Team. R: A language and environment for statistical computing. Vienna, Austria: R Foundation for Statistical Computing; 2025. Available: http://www.R-project.org/

